# Bilateral asymmetric hip stiffness applied by a robotic hip exoskeleton elicits kinematic and kinetic adaptation

**DOI:** 10.1101/2023.02.06.527337

**Authors:** Banu Abdikadirova, Mark Price, Jonaz Moreno Jaramillo, Wouter Hoogkamer, Meghan E. Huber

## Abstract

Wearable robotic exoskeletons hold great promise for gait rehabilitation as portable, accessible tools. However, a better understanding of the potential for exoskeletons to elicit neural adaptation—a critical component of neurological gait rehabilitation—is needed. In this study, we investigated whether humans adapt to bilateral asymmetric stiffness perturbations applied by a hip exoskeleton, taking inspiration from asymmetry augmentation strategies used in split-belt treadmill training. During walking, we applied torques about the hip joints to repel the thigh away from a neutral position on the left side and attract the thigh toward a neutral position on the right side. Six participants performed an adaptation walking trial on a treadmill while wearing the exoskeleton. The exoskeleton elicited time-varying changes and aftereffects in step length and propulsive/braking ground reaction forces, indicating behavioral signatures of neural adaptation. These responses resemble typical responses to split-belt treadmill training, suggesting that the proposed intervention with a robotic hip exoskeleton may be an effective approach to (re)training symmetric gait.

## I. INTRODUCTION

Impaired gait is a common symptom of neurological disorders that can restrict one’s mobility, negatively impacting their quality of life. Such gait impairments include slower gait speed, reduced step length, and asymmetric gait kinematics [1], [2]. Because the excessive, abnormal joint loading caused by asymmetric gait can also degrade musculoskeletal health [3], interventions that restore symmetric gait patterns are greatly needed.

Split-belt treadmill training is a promising approach to correct for gait asymmetry caused by neurological disorder [4]. In split-belt treadmill training, asymmetries in step length are induced by running the belts of a split-belt treadmill at different speeds [4]. Prior results show that split-belt treadmill training elicits neural adaptation and improves step-length symmetry in post-stroke individuals, but this improvement only partially transfers to overground walking [5], even with repetitive training [6]. Limitations in transfer have also been observed in load-based measures critical to rehabilitation outcomes such as weight bearing or propulsion [7], [8]. Furthermore, access to split-belt treadmill training is limited by proximity to a facility with the resources to house and operate the technology, as well as the frequency of availability to an individual patient. Wearable robotic technology may offer a solution to overcoming the current limitations of split-belt treadmill training.

In general, wearable robotic exoskeletons hold great promise for gait rehabilitation. The development of new actuators optimized for working in concert with musculoskeletal dynamics [9], [10] as well as innovations in exoskeleton control [11] have enabled the design of wearable devices that can augment human performance (i.e., reduce the metabolic cost of walking [12]) and assist impaired mobility (i.e., reduce crouching in children with cerebral palsy [13]).

Compared to split-belt treadmills, wearable robotic exoskeletons are more accessible due to their smaller size, lower cost, and easier usability during activities of daily living. Also, wearable robots can potentially induce changes not induced by split-belt treadmills, as they apply torques directly onto the lower limb joints as opposed to applying speed constraints to the feet. Whether the same neural adaptation capable of improving gait symmetry in patients from split-belt training can also be induced by a robotic exoskeleton is still an open question.

In prior work, we identified behavioral signatures of neural adaptation to unilateral asymmetric stiffness applied with a hip exoskeleton [14]. Specifically, returns to baseline behavior and post-intervention after-effects were observed in spatiotemporal measures of gait. However, participants did not fully converge towards adopting symmetric gait patterns as expected. One possible explanation for this result is that the gait asymmetry created by applying unilateral stiffness may not have been large enough to be recognized as an error to be corrected by the nervous system.

The purpose of the present study was to investigate whether applying a bilateral asymmetric stiffness using a hip exoskeleton elicits neural adaptation towards symmetric gait. Instead of only applying a positive stiffness to one hip joint as in prior work [14], a positive stiffness was applied to one hip joint and a negative stiffness on the other to a induce larger spatial gait asymmetry (Fig. 1A-C). We additionally investigated the vertical and anterior-posterior ground reaction forces to determine if our exoskeleton intervention caused changes in weight-bearing or propulsive/braking force symmetry.

**Fig. 1.**
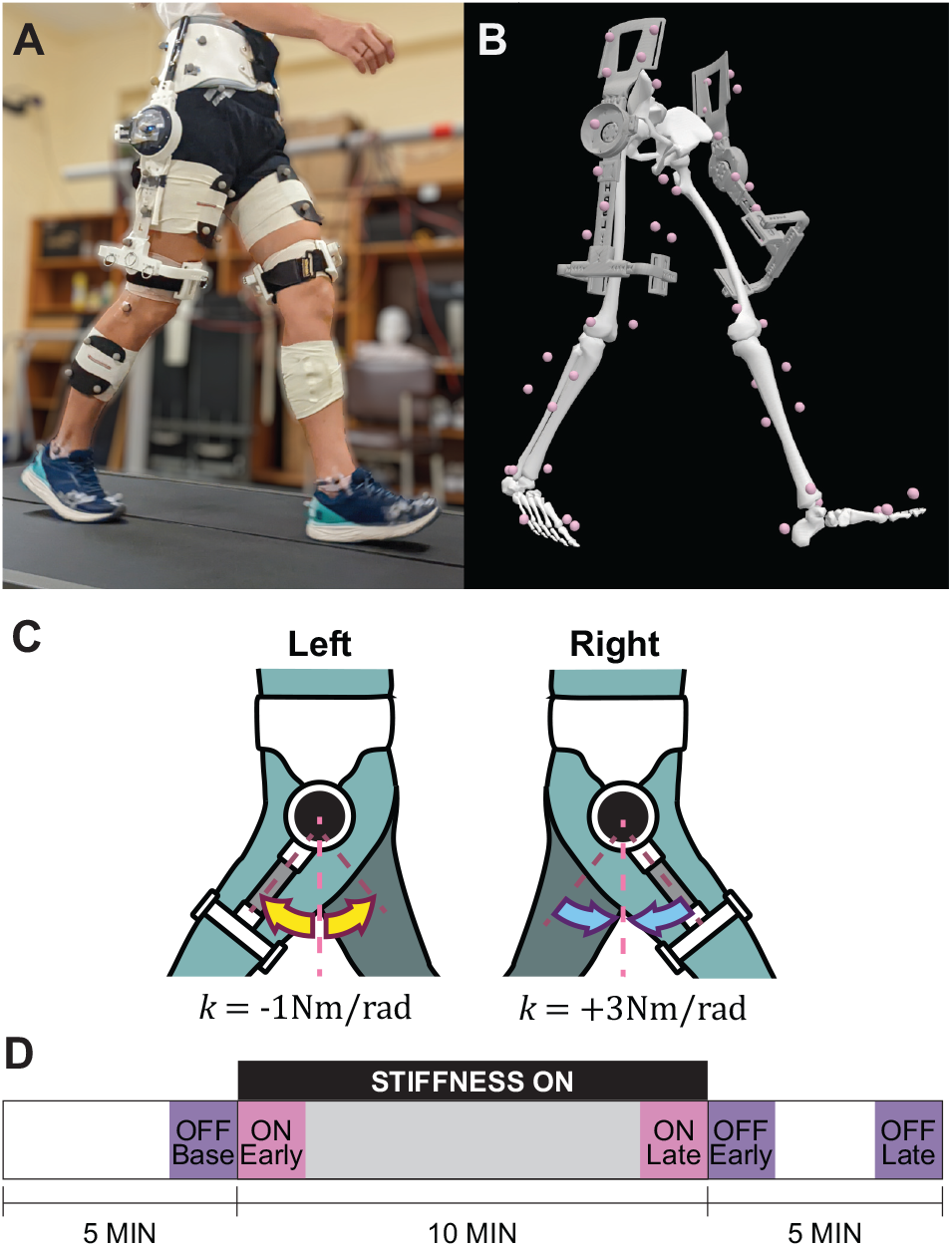
**A** Participants walked on an instrumented treadmill wearing a bilateral hip exoskeleton and reflective markers for motion capture. **B** Lower limb and exoskeleton joint kinematics were calculated by converting motion capture data into individualized models of each participant in OpenSim. **C** The exoskeleton exerted attractive and repulsive torques expressed as positive and negative stiffness on the right and left sides, respectively. **D** Experimental protocol consisted of 5 minutes of baseline walking (stiffness controller OFF), 10 minutes of walking with stiffness ON, and 5 minutes of walking with stiffness OFF.

The remainder of the paper is organized as follows.

Section II describes the experimental design and methods used to evaluate the effect of bilateral modulation of hip joint stiffness on gait kinematics and asymmetry. Section III presents results which are discussed in Section IV. Concluding remarks follow in Section V.

## II. METHODS

### A. Participants

A total of six healthy, young adults (gender: one female, five male; age: 24.7 ± 3.6 years; height: 1.75 ± 0.09 m; mass: 75.2 ± 18.8 kg) took part in this study. None had previously worn a hip exoskeleton nor partook in a similar experiment. All subjects signed informed written consent before the experiment. The experimental protocol was reviewed and approved by the Institutional Review Board of the University of Massachusetts Amherst.

### B. HRSL Hip Exoskeleton

The hip exoskeleton used in this study was a bilateral configuration of the exoskeleton used in [14] (Fig. 1A). In this configuration, the mass of the exoskeleton is approximately 3 kg. The exoskeleton is suspended over the hips by an adjustable waist harness, and custom frames are fitted to front and back of the thighs via adjustable pin-lock mechanisms. Torque is transmitted about the flexion/extension axis of each hip joint through the trunk and thighs. Passive hinges allow for hip abduction and adduction in the frontal plane.

Each actuator contains a brushless DC motor with a 6:1 gearhead and an absolute encoder, along with additional sensors and electronics (ActPack 4.1, Dephy, Maynard, MA, USA). Output torque from the actuator is estimated and controlled by sensing electrical current in the motor. High-level control and operation is handled through a Raspberry Pi 4 (Raspberry Pi Ltd, Cambridge, UK) microcomputer. Because the current experiment was performed on a treadmill, the power source and microcomputer were located offboard.

### C. Stiffness Controller

For this experiment, the actuators emulated virtual, torsional springs using the following control law: *τ* = *kθ*, where *τ* is the motor torque applied to the hip joint, *θ* is the hip angle defined relative to an upright standing position as measured by the encoder in the actuator, and *k* is the stiffness of the spring.

The stiffness value can be configured to be either positive or negative. For positive stiffness, the exoskeleton applies a torque about the hip joint such that it pulls the leg towards the upright standing position. For negative stiffness, the exoskeleton applies a torque around the hip joint to repel the leg away from the upright standing position (Fig. 1C). For this experiment, the stiffness of the spring was set to 3.0 Nm/rad and −1.0 Nm/rad on the right and left sides of the exoskeleton, respectively. The stiffness controller was turned off by setting *τ* = 0.

### D. Experimental Procedure

The experiment took place in a single session for each participant. Data collection was preceded by fitting the exoskeleton to the participant’s waist and thighs until snug. The exoskeleton was then removed and the participant was recorded standing in a calibration pose before walking for 2 minutes on the treadmill to obtain a baseline of their normal walking biomechanics. The participant then re-donned the exoskeleton and was recorded in another standing calibration pose to ensure accurate model configuration with and without the exoskeleton. Each participant then performed one trial in which they walked at 1.30 m/s for a total of 20 minutes on an instrumented split-belt treadmill (Bertec Corporation, Columbus, OH, USA). The stiffness controller was turned off during the first five minutes, turned on for ten minutes, and turned off again for the remaining five minutes (Fig. 1D).

### E. Data Collection

A total of 52 reflective markers were placed on each participant for model scaling and motion tracking. Marker positions were recorded at 100Hz with an eight-camera motion capture system (Qualisys, Inc., Gothenburg, Sweden). Ground reaction forces were recorded at 1000Hz with a pair of force plates located under the treadmill belts (Bertec Corporation, Columbus, OH, USA). Markers were placed to locate the pelvis, thighs, shanks, feet, and all segments of the bilateral exoskeleton in 3D space. The motion capture system was calibrated immediately before each experiment per manufacturer specifications.

### F. Data Processing

3D models of the human-exoskeleton system were created in OpenSim 4.3 [15] for each participant (Fig. 1B). The exoskeleton is represented by two three-segment systems, each consisting of a waist harness segment, a motor segment, and a thigh frame segment. The segments are connected by two revolute joints: an ab/adduction hinge connecting the motor to the waist harness, and a flexion/extension joint between the motor and thigh frame representing the motor angle. The exoskeleton systems are not constrained relative to the human skeleton or to each other; each waist harness segment is connected to the pelvis by a six degree-of-freedom joint, and the thigh segment is not connected to the human thigh segment at all. This configuration allows the exoskeleton and human joint kinematics to be quantified independently. This method builds on our earlier work in [14] creating accurate 3D models with a unilateral hip exoskeleton.

Marker positions were filtered with a fourth-order zero-lag Butterworth low-pass filter (6 Hz) using the filtfilt function in MATLAB (The Mathworks, Natick, MA) to remove high frequency noise. Marker positions captured during quiet standing were used to scale OpenSim models and place virtual markers for each participant. Joint kinematics were calculated by using a global least-squares optimization which minimizes weighted model marker position errors relative to experimental marker positions, subject to model joint constraints [16].

### G. Dependent Measures

The gait cycle was defined as beginning at heel-strike of the right leg (0%) and concluding at the following heelstrike of the same leg (100%). Heel-strike was determined as occurring during the rising edge of the right vertical ground reaction force (GRF) through a threshold of 10 N. The joint kinematics and GRFs from each trial were segmented to calculate the following dependent measures for each stride.

1. *Spatiotemporal measures:* The spatial aspects of gait were quantified by the angular range of motion (RoM) of the left and right hip joints, and the step lengths of the left and right legs. Step length was defined as the anterior-posterior distance between heel markers at heel-strike of the respective leg. The temporal aspect of each stride was characterized by step time for each limb. Step time was defined as the time between heel-strike and the following heel-strike of the opposite limb.
2. *Kinetic measures:* Load-bearing characteristics for each stride were quantified by the peak propulsive, braking, and vertical GRF for each limb, scaled by the mass of the participant.
3. *Asymmetry definition:* The asymmetry between left and right sides was quantified by calculating a ratio defined as

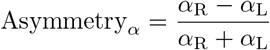

where *α* represents the dependent measure for the right (R) and left (L) leg.

### H. Statistical Analysis

For each participant, the mean of each dependent measure was calculated during each of the following conditions: the terminal 10 strides in the baseline phase with the stiffness controller off (OFF-Base), the initial (ON-Early) and terminal (ON-Late) 10 strides in the exposure phase with the stiffness controller on, and the initial (OFF-Early) and terminal (OFF-Late) 10 strides in the post-exposure phase with the stiffness controller off. These conditions are summarized in Fig. 1C.

One-way repeated measures analysis of variances (ANOVAs) were conducted to assess the effect of condition (OFF-Base, ON-Early, ON-Late, OFF-Early, OFF-Late) on each of the dependent measures. If Mauchly’s test of sphericity was statistically significant (i.e., the assumption of equal variances of the differences between all combinations of conditions was violated), the Greenhouse-Geisser correction factor was applied to the degrees of freedom of the ANOVA.

A significant effect of condition was followed up with planned comparisons in the form of pairwise t-tests between the following conditions: (1) OFF-Base vs. ON-Early, (2) OFF-Base vs. ON-Late, (3) OFF-Base vs. OFF-Early, (4) OFF-Base vs. OFF-Late. A Bonferroni correction was applied to the reported p-values to control for Type I errors across these four comparisons. These planned comparisons were conducted to assess behavioral signatures of neural adaptation [17].

The statistical analyses were performed using a custom script in MATLAB (MathWorks, Natick, MA). For all statistical tests, the significance level was set to *α* = 0.05.

## III. RESULTS

Fig 2 illustrates how the dependent measures change over strides and across conditions for a representative participant. The ANOVAs revealed a statistically significant effect of condition on all spatio-temporal and kinetic gait measures. Detailed results are discussed in the section below. The results of planned comparisons are summarized in Fig. 3 and Fig. 4 for the spatiotemporal and kinetic measures, respectively.

**Fig. 2.**
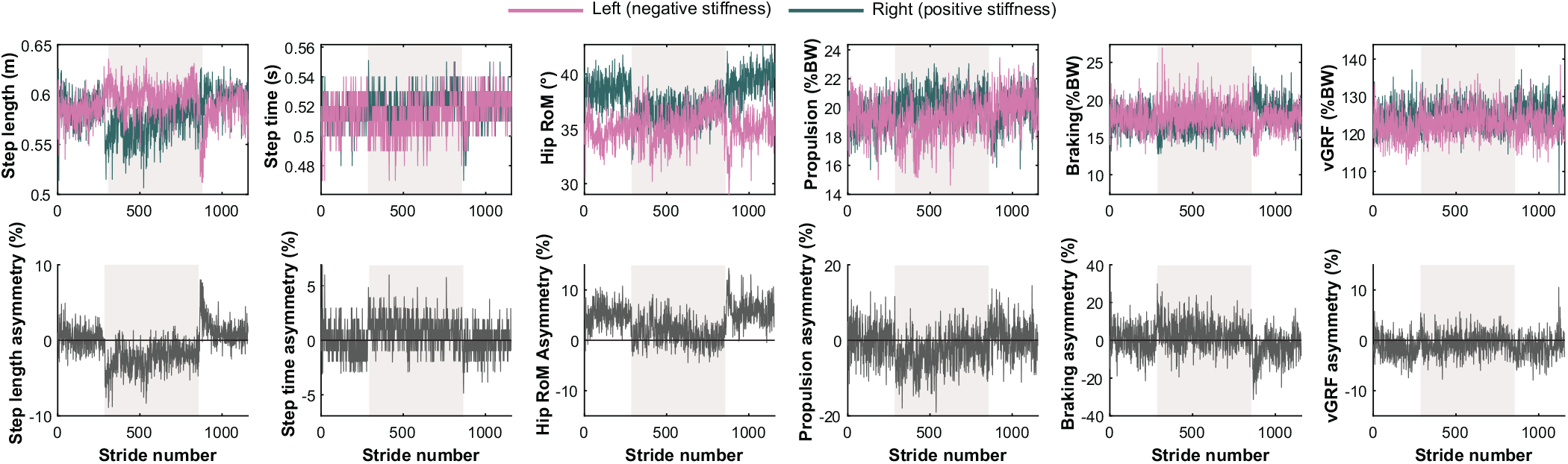
Dependent measures of a representative participant. Shaded regions represent when the stiffness controller was on.

**Fig. 3.**
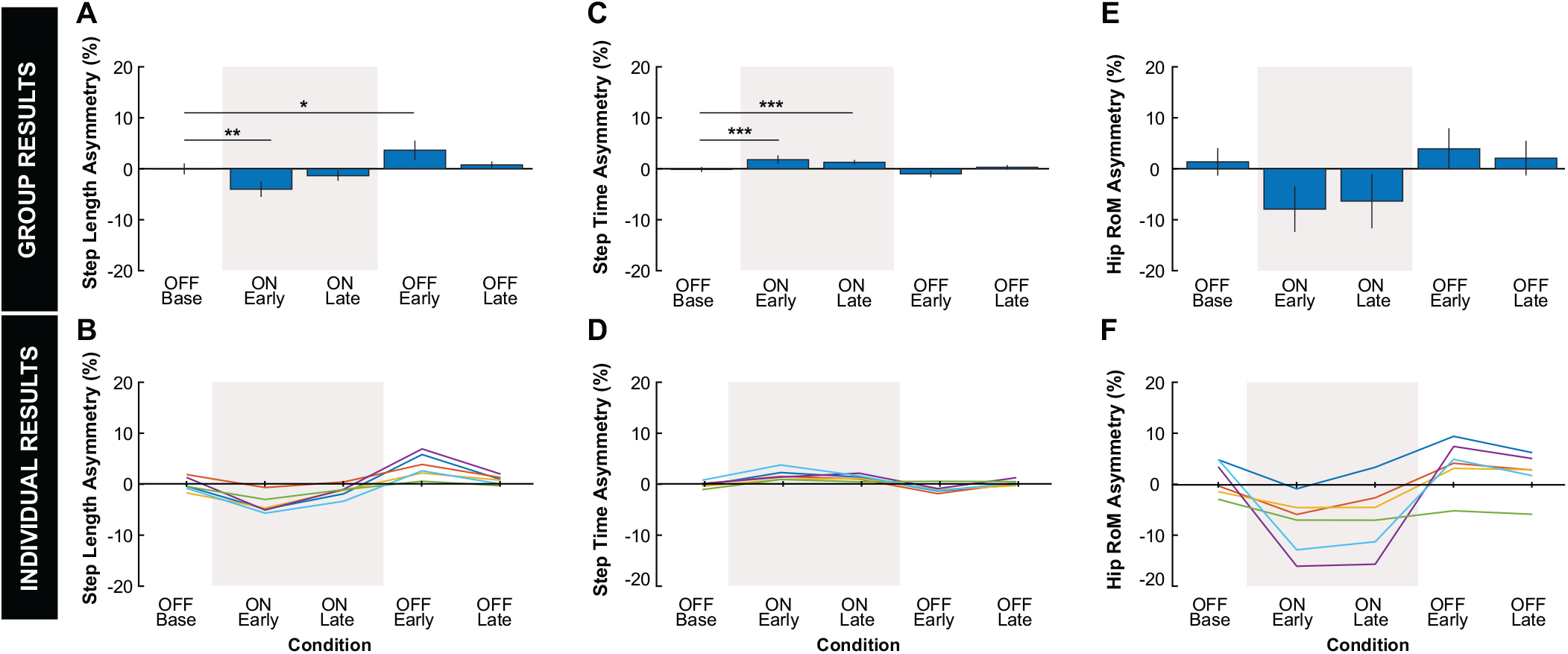
Spatiotemporal results. **A:** Group average and **B:** individual results for step length asymmetry. **C:** Group average and **D:** individual results for step time asymmetry. **E:** Group average and **F:** individual results for hip RoM asymmetry. **A, C, E:** Error bars represent two standard errors of the mean. Shaded regions represent when the stiffness controller was on. The ANOVAs found a statistically significant effect of condition on all spatiotemporal measures. *, **, and *** indicate that the planned comparison between conditions was statistically significant with *p* < 0.05, *p* < 0.01, *p* < 0.005, *p* < 0.001, respectively. **B, D, F:** Color indicates the different individual subjects.

**Fig. 4.**
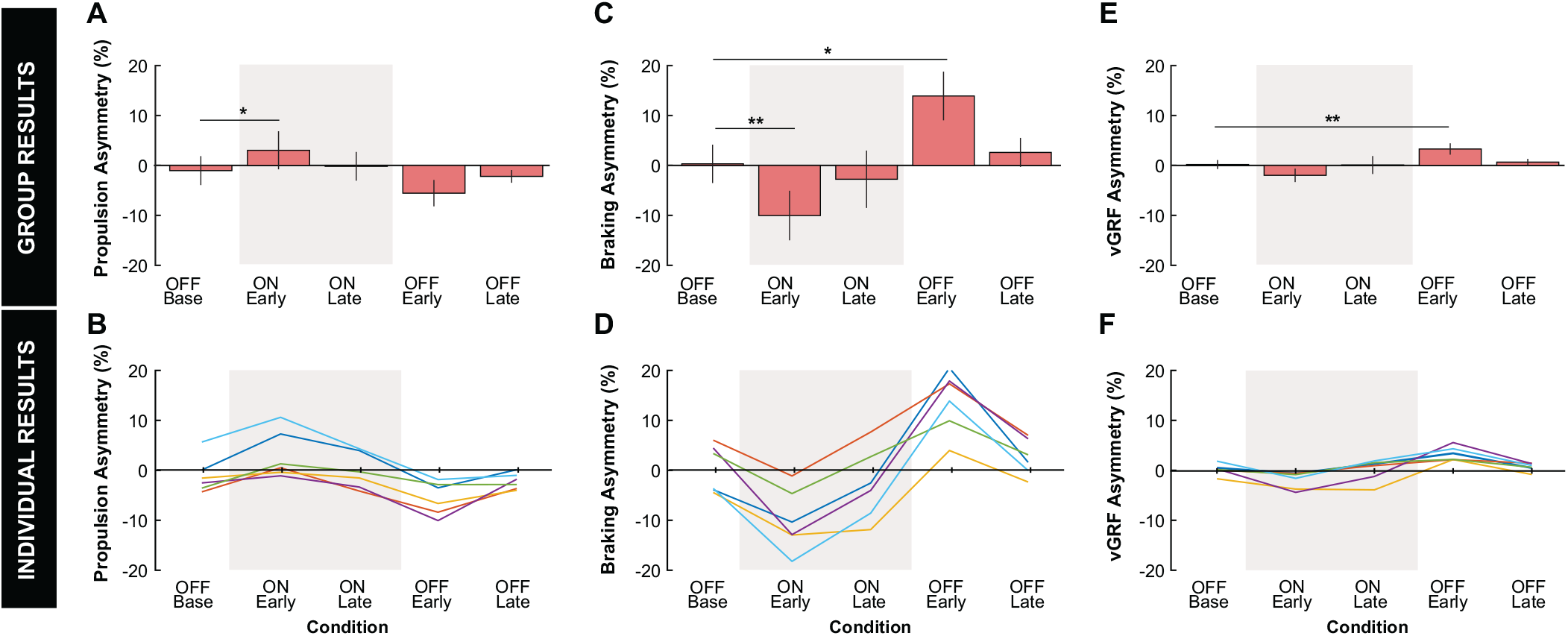
Kinetic results. **A:** Group average and **B:** individual results for propulsion asymmetry. **C:** Group average and **D:** individual results for braking asymmetry. **E:** Group average and **F:** individual results for vertical GRF (vGRF) asymmetry. **A, C, E:** Error bars represent two standard errors of the mean. Shaded regions represent when the stiffness controller was on. The ANOVAs found a statistically significant effect of condition on all spatiotemporal measures. *, **, and *** indicate that the planned comparison between conditions was statistically significant with *p* < 0.05, *p* < 0.01, *p* < 0.005, *p* < 0.001, respectively. **B, D, F:** Color indicates the different individual subjects.

### A. Spatiotemporal Measures

#### 1) Step length

The ANOVA revealed a statistically significant effect of condition on Step Length asymmetry (*F*_4,20_ = 26.48, *p* < .001, Fig. 3A). During the OFF-Base condition, step length was symmetric (*M* = −0.04%). Turning on the stiffness controller induced an asymmetry in the negative direction as seen in the ON-Early condition (*M* = −4.02%). Over time, the magnitude of negative asymmetry reduced such that there was no statistical difference between the ON-Late and OFF-Base conditions. Turning off the stiffness controller induced a positive asymmetry (*M* = 3.64%) as seen in the OFF-Early condition, but again this asymmetry was reduced such that there was no statistical difference between the OFF-Late and OFF-Base conditions.

#### 2) Step time

The ANOVA revealed that the effect of condition on step time asymmetry was statistically significant (*F*_4,20_ = 13.87, *p* < .001, Fig. 3C). During the OFF-Base condition, step time was symmetric (*M* = −0.14%). Turning on the stiffness controller induced a positive asymmetry in the ON-Early condition (*M* = 1.79%), which was maintained even until the ON-Late condition (*M* = 1.27%). When the stiffness controller was turned off, step time asymmetry in neither the OFF-Early condition nor the OFF-Late condition were statistically different from the OFF-Base condition.

#### 3) Hip RoM

The ANOVA revealed that the effect of condition on Hip RoM asymmetry was statistically significant (*F*_1.25,6.27_ = 9.63, *p* = .017, Fig. 3E). However, due to high within-participant variance, none of the planned comparisons for each condition with respect to baseline reached statistical significance with the Bonferroni correction applied.

### B. Kinetic Measures

#### 1) Propulsive ground reaction forces

The ANOVA revealed a statistically significant effect of condition on peak propulsion asymmetry (*F*_4,20_ = 15.74, *p* < .001, Fig. 4A). During the OFF-Base condition, the peak propulsion GRFs were symmetric (*M* = −1.03%). Turning on the stiffness controller induced a positive asymmetry in the ON-Early condition (*M* = 3.0%), which disappeared by the ON-Late condition. A marginal aftereffect in peak propulsion was observed, but was not significant with the Bonferroni correction applied (*M* = −5.5%). This after effect washed out to OFF-Base values by the OFF-Late condition.

#### 2) Braking ground reaction forces

The effect of condition on peak braking asymmetry was statistically significant (*F*_1.87,9.33_ = 32.77, *p* < .001, Fig. 4C). During the OFF-Base condition, the peak braking GRFs were symmetric (*M* = 0.32%). Opposite of propulsion, turning on the stiffness controller induced a negative asymmetry in the ON-Early condition (*M* = −10.0%), the magnitude of which reduced to OFF-Base values by ON-Late condition. An aftereffect was observed, where a positive peak braking asymmetry was observed in the OFF-Early condition (*M* = 13.9%) during OFF-Early. This aftereffect washed out to OFF-Base values by the OFF-Late conditions.

#### 3) Vertical ground reaction forces (vGRFs)

The effect of condition on peak vGRF asymmetry was statistically significant (*F*_4,20_ = 14.17, *p* < .001, Fig. 4E). During the OFF-Base condition, the peak vGRFs were symmetric (*M* = 0.2%). Turning on the stiffness controller did not induce a statistically significant asymmetry compared to OFF-Base values in neither the ON-Early nor ON-Late conditions. An aftereffect was observed, with peak vGRF a positive asymmetry observed during OFF-Early condition (*M* = 3.3%). This aftereffect washed out to OFF-Base values by the OFF-Late condition.

## IV. DISCUSSION

### A. Behavioral Comparisons in Unilateral vs. Bilateral Asymmetry Intervention

Compared to our previous unilateral stiffness controller [14], the bilateral configuration of the exoskeleton induced larger immediate changes in step length asymmetry. From 3A, the initial asymmetry in step length was ~ −4% during ON-Early condition, while the unilateral configuration only induced an asymmetry of ~ −2%. With both unilateral and bilateral stiffness configurations, the magnitude of the step asymmetry reduced as participants continued to walk with the respective stiffness controllers on. The bilateral configuration also resulted in more obvious aftereffect in step length asymmetry (~4%) compared to the unilateral configuration (~2%). However, these aftereffects washed out in both stiffness configurations. For the bilateral stiffness controller, the effect of condition on step time asymmetry was weaker compared to the spatial dependent measures. However, it still induced larger changes in step time asymmetry compared to a unilateral stiffness configuration. Turning the stiffness controller induced an asymmetry of ~2% in the bilateral configuration compared to ~1% in the unilateral configuration. In both configurations, an aftereffect in step time asymmetry was not observed.

In our earlier work [14], the unilateral configuration induced an asymmetry in all spatiotemporal parameters, even when the stiffness controller was off. In this study, the bilateral configuration of the exoskeleton eliminated that baseline asymmetry, which could be due to either more even distribution of the weight of the exoskeleton on the body and/or a more symmetrical imposition of movement constraints caused by the exoskeleton geometry.

### B. Comparison to Split-Belt Treadmill Adaptation

Typical hallmarks of adaptation during split-belt treadmill training with healthy participants include (1) a return to baseline behavior over time during the intervention and (2) an aftereffect upon cessation of the intervention in measures of step length asymmetry and ground reaction forces. The results of this study showed that similar effects can be elicited with a wearable exoskeleton acting locally on the hip joints.

As with split-belt treadmills, the clearest signs of adaptation occurred with step length [4] and braking GRF magnitude [18]-[21]. It is likely that these measures are related, as pendular walking mechanics dictate that the braking component of the resultant GRF at heel strike depends on the limb angle relative to the treadmill belt [22].

Also similar to split belt treadmill studies, we observed a weaker response in propulsion than braking. Propulsion aftereffects have been observed in some split-belt treadmill walking experiments [19], [20] and marginally evidenced or not observed in others [18], [21]. Our results suggest that a propulsion aftereffect may be present but its statistical significance is marginal. The exoskeleton may cause neuromotor adaptations in propulsion asymmetry for two reasons that are not possible with a split-belt treadmill alone: (1) The exoskeleton can directly influence hip extension, thereby influencing the extension angle of the limb at push-off, which is strongly associated with propulsion [7], and (2) the exoskeleton produces hip torques during push-off, which may either reduce the amount of push-off force transmitted via plantarflexion torque needed to initiate swing on the negative stiffness side, or increase it on the positive stiffness side.

Interestingly, we found that participants adapted measures that require endpoint control (step length, foot-ground interaction forces) toward symmetry when the stiffness controller was active. However, they maintained prolonged local asymmetry (hip RoM) where the exoskeleton directly applied torques. A possible explanation is that symmetry in endpoint or global measures are prioritized by the nervous system. For instance, if joint-level symmetry is energetically costly or uncomfortable, the neuromotor system might tolerate joint-level asymmetry if global symmetry can be achieved without it. This factor may be responsible for the resemblance of these adaptation patterns to those elicited by split-belt treadmills, which interact directly with the feet (i.e. the endpoints of the limbs) during stance. However, this speculation remains to be tested.

### C. Limitations and Future Work

Though our results showed important findings about how humans adapt to bilateral asymmetrical stiffness applied by a hip exoskeleton, the experiment was conducted with a small and uniform population of subjects. It is likely that we were unable to observe statistical significance for some measures with seemingly large differences in the mean values due to a lack of statistical power. Future experiments will be conducted with larger and more diverse samples to test whether their response to a bilateral stiffness application is consistent among various groups.

Our controller allowed only a limited range of stiffness configurations. For instance, positive stiffness values above 3 Nm/rad resulted in less stable controller performance in the asymmetric bilateral configuration. Higher positive stiffness values could potentially help to exaggerate walking asymmetry even more and elicit a stronger adaptation response. However, our results also show that relatively mild torques applied to the hip joint may induce gait asymmetry and induce behavioral signatures of neural adaptation.

Our hip exoskeleton was designed to be highly adjustable to fit a larger population of people of different sizes. Despite this, we did not quantify exoskeleton fit, which varied between participants. Misalignment between the motors and the hip joints could have been introduced due to sagging of the exoskeleton over time, which might have altered the effect of the exoskeleton on the gait.

Despite these limitations, the ability for a hip exoskeleton to reproduce split belt treadmill adaptation behaviors is extremely promising. Our future work includes the study of similar interventions overground to determine if the effects can be reproduced, and critically, improve transference of the effects to unassisted overground walking compared to split belt treadmill training. Furthermore, while our experiment was conducted on an instrumented dual-belt treadmill, the belts were run at the same speeds for all conditions, meaning that this intervention can be carried out effectively on an inexpensive, single-belt treadmill. Finally, we expect to study the effect of these interventions on clinical populations if our studies with healthy participants continue to produce promising results.

## V. CONCLUSIONS

This study investigated the changes in spatiotemporal and kinetic gait measures by applying asymmetric bilateral stiffness at the hip joints with the HRSL robotic hip exoskeleton. The results indicated stronger behavioral signatures of neural adaptation compared to a unilateral stiffness configuration. As in split-belt treadmill studies, turning the stiffness controller on initially induced asymmetry. Over time, while the exoskeleton was still on, subjects tended towards symmetrical step lengths and propulsive/braking ground reaction forces. This indicates that the nervous system may be treating these asymmetries as errors that need to be corrected. Turning the stiffness controller off resulted in aftereffects which were in an opposing direction to the initially induced gait asymmetries. Even though the aftereffects of applying bilateral stiffness around the hip joints were not long lasting, observed signs of neural adaptation suggest that the proposed intervention has potential as an effective gait neurorehabilitation technique.

## Acknowledgement

We would like to thank Savannah Macero and Robert Bennett for their assistance during pilot tests for this protocol. We would also like to thank Herlandt Lino, Savannah Macero, and Keanan Bradley for their assistance processing motion capture data for this study.

